# Knockdown of fibrillarin reveals cell- and virus-dependent restriction of henipavirus replication and enhanced antiviral immune signalling

**DOI:** 10.64898/2026.11.02.763686

**Authors:** Rasan Mohamed Sathiqu, Christina L. Rootes, Aaron M. Brice, Alessandro T. Caputo, Sinead M. Williams, Carlos H. M. Rodrigues, Ryan J. Farr, Marina R. Alexander, Cameron R. Stewart

**Author notes:** Correspondence: Cameron Stewart.

## Abstract

The *henipavirus* genus includes several highly pathogenic zoonotic viruses that lack medical countermeasures. The 2’-O-methyltransferase fibrillarin (FBL) has been identified by genome-wide functional genomics studies as a henipavirus pro-viral factor. While FBL is a potential target for host-directed antiviral therapeutics, its impact on henipavirus replication in cell types relevant to henipavirus disease has not been explored. Here we demonstrate that depleting cellular FBL using short interfering RNAs (siRNAs) has varying impacts on henipavirus replication, depending on cell type and virus. FBL depletion in small airway epithelial cells (HSAEC1-KT) significantly impaired viral gene transcription for Hendra virus (HeV) genotypes 1 and 2 (HeV-g1, HeV-g2), and Nipah virus (NiV) Bangladesh strain (NiV-B), but not NiV Malaysia (NiV-M) strain or Cedar virus (CedV). Conversely, reduced FBL expression in bronchial epithelial (NuLi-1) cells, aortic endothelial (Telo HAEC) cells or neuroblastoma (BE(2)-M17) cells did not inhibit HeV-g1 replication. Transcriptomic analyses revealed that impaired virus replication in FBL knockdown cells was associated with enhanced activation of innate immune pathways and robust upregulation of interferon-stimulated genes (ISGs). Together, these findings demonstrate a virus- and cell-type dependent impact of FBL on henipavirus replication and support a role for FBL in modulation of the antiviral innate immune response.

## INTRODUCTION

The *Henipavirus* genus, belonging to the *Paramyxovirus* family, contains several zoonotic, negative-sense RNA viruses, including Hendra virus (HeV), Nipah virus (NiV), Cedar virus (CedV), Ghana virus and Angavokely virus. To date HeV and NiV are the only members known to cause human disease, both with high case fatality rates. HeV was first identified during an outbreak in Queensland, Australia in 1994 [1], followed by the identification of NiV in Malaysia in 1998 [2]. Since its discovery, NiV has been detected across South and Southeast Asia, with near-yearly outbreaks reported across Malaysia, Singapore, Bangladesh, India, and the Philippines [3]. In contrast, HeV remains endemic to Australia and has resulted in sporadic, localised outbreaks. Both HeV and NiV are maintained in nature by *Pteropus* species fruit bats, which serve as their natural reservoirs [4,5]. Spillover can occur through transmission from bats to intermediate amplifying hosts, horses for HeV and pigs for NiV before infecting humans. HeV and NiV infections in humans are characterised by severe respiratory disease and encephalitis, often progressing rapidly [1,2,6,7]. Of particular concern is the NiV-Bangladesh (NiV-B) strain, which causes almost annual outbreaks in Bangladesh and India and is associated with case fatality rates frequently exceeding 70%, as well as documented human-to-human transmission [8].

In recognition of this threat, in July 2024 the World Health Organization updated its list of priority pathogens to include henipaviruses as agents capable of causing a future viral pandemic [9]. This prioritisation aims to accelerate research efforts focused on understanding henipavirus biology and developing effective medical countermeasures. At present, therapeutic options are limited, with none approved for use in humans and a single licensed subunit vaccine for use in horses targeting the HeV glycoprotein (G) [10]. Several experimental human countermeasures are under development, including vaccines and monoclonal antibodies targeting the viral attachment (G) and fusion (F) glycoproteins [11–13]. Monoclonal antibodies directed against the G protein have demonstrated potent neutralising activity in preclinical animal models [14] and have been used on a compassionate basis in a small number of human exposures [15]. The growing diversity of characterised henipaviruses, coupled with their high mortality, highlights the need for broadly effective antiviral strategies.

As with all viruses, henipaviruses rely extensively on host cellular factors for infection and replication. A genome-wide functional genomics screen identified FBL, a conserved nucleolar 2’-O-methyltransferase involved in ribosomal RNA processing and ribosome biogenesis, as a critical host factor for HeV and NiV replication in immortalised cervical cancer (HeLa) cells [16]. FBL was shown to be required during the replication stage of the henipavirus infection cycle, with its catalytic activity essential for viral RNA synthesis and viral protein production [16], although the precise mechanistic role remains unclear. Depletion of FBL was also shown to impact other medically-relevant paramyxovirus genera, namely the morbillivirus, measles virus, and the orthorubulavirus, mumps virus. The pan-viral activity of FBL across the paramyxovirus family makes it an attractive target for the development of host-directed antiviral (HDA) strategies.

In this study, we investigate the impact of inhibiting FBL on the replication kinetics of different henipaviruses in cell types relevant to henipavirus tropism. HSAEC1-KT and NuLi-1 cells model small airway and bronchial epithelial cells, respectively, reflecting the respiratory epithelium as an important initial site of viral entry, replication and shedding [17,18]. TeloHAEC cells represent human vascular endothelial cells, a major target of henipaviruses during systemic infection, where viral replication contributes to vasculitis and vascular leakage [17,19]. BE(2)-M17 neuroblastoma cells serve as a surrogate for neuronal cells, capturing the neurotropism of HeV and NiV and their capacity to invade the central nervous system [19]. Unlike HeLa cells, which are highly transformed cervical carcinoma cells, HSAEC1-KT, NuLi-1 and TeloHAEC cells are immortalised derivatives of normal human tissues that retain characteristics of their primary counterparts, while BE(2)-M17 cells are a neuroblastoma-derived surrogate neuronal model. Together, these cell types enable examination of host–virus interactions across the principal tissues associated with henipavirus disease in humans. In addition to evaluating antiviral efficacy of siRNA knockdown of host genes, we have employed RNAseq to identify potential mechanisms for host factor-dependent virus replication.

## RESULTS

### Characterising HeV-g1 infection permissiveness and FBL knockdown across cell types

We firstly established *in vitro* systems for cells receptive to both henipavirus infection and siRNA-mediated gene knockdown. HeV-g1 replication was assessed in four cell lines representing endothelial (TeloHAEC), lung small airway epithelial (HSAEC1-KT), lung bronchial epithelial (NuLi-1) and neuronal (BE(2)-M17) origins. Cells were infected at cell line-specific multiplicities of infection (MOIs), and viral replication was quantified at 1 h (inoculation levels) and 24 h post-infection (hpi) by measurement of HeV *N* gene expression (Fig. 1A) and infectious virus titres (Fig. 1B). Across all cell types, HeV *N* gene expression and infectious virus were detected at inoculation levels and increased at 24 hpi. BE(2)-M17 cells showed the highest increases in viral mRNA (∼730,000-fold over inoculum), followed by HSAEC1-KT (32,000-fold) NuLi-1 (19,500-fold), then TeloHAEC (1,000-fold). In contrast, the highest increase in virus titres were observed in HSAEC1-KT cells and BE(2)-M17 cells (2.31 log_10T_CID_50 i_ncrease), followed by NuLi-1 cells (2.28 log_10T_CID_50 i_ncrease). Telo HAEC cells demonstrated a very low increases in virus titre (0.69 log_10T_CID_50)_.To assess susceptibility of the cell line panel to siRNA-mediated knockdown of FBL, each cell line was transfected with 40 nM siNEG or siFBL. Reduced FBL protein expression was observed in after 120 h in TeloHAEC and HSAEC1-KT cells and at 72 h in NuLi-1 and BE(2)-M17 cells (Fig. 1C).

**Figure 1:**
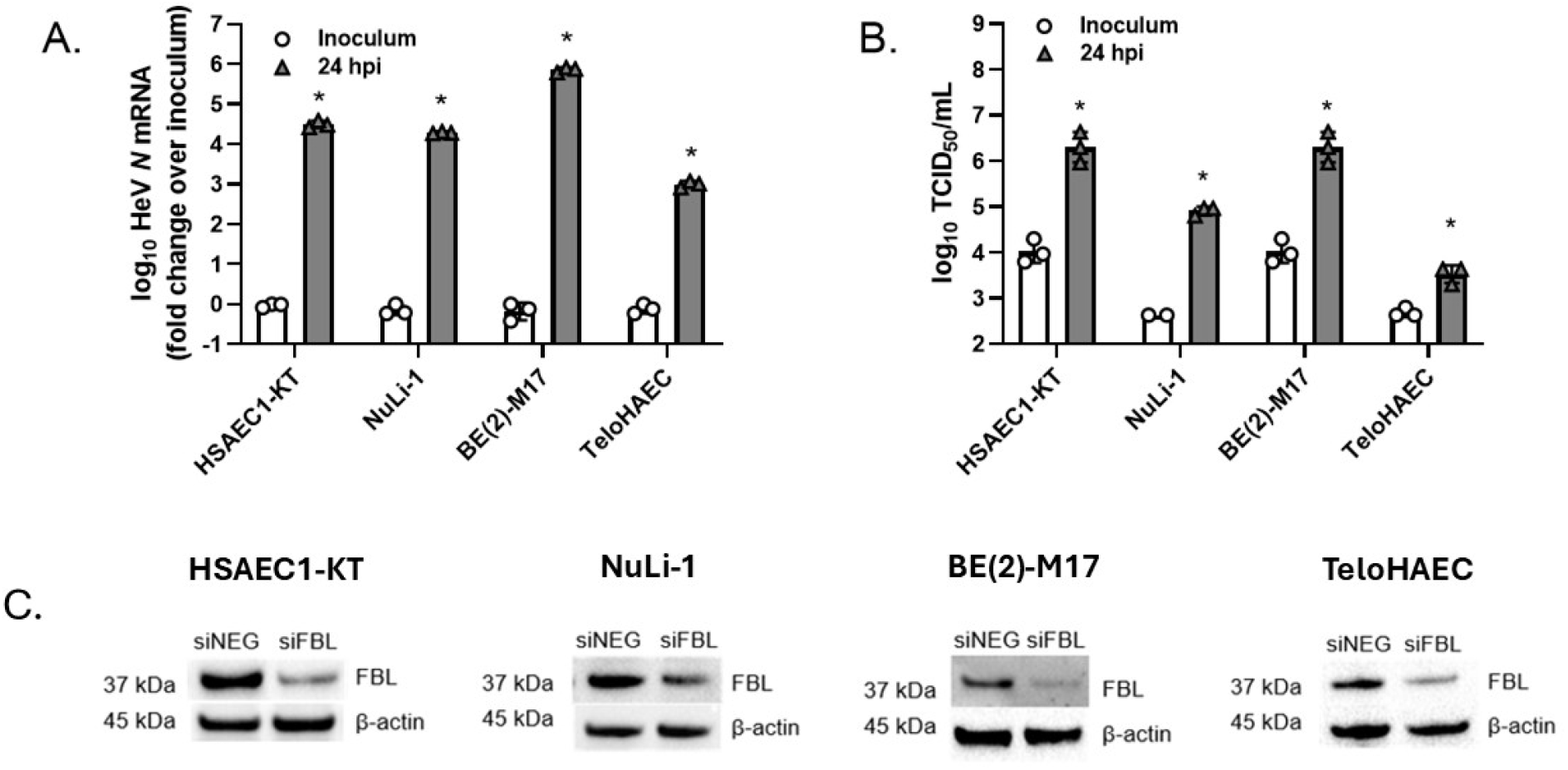
Growth kinetics of HeV-g1 in small airway epithelial, bronchial epithelial, aortic endothelial and neuronal-like neuroblastoma cells. Infection permissiveness in HSAEC1-KT cells (MOI=0.2), NuLi-1 cells (MOI=1), BE(2)-M17 cells (MOI=0.05) and TeloHAEC cells (MOI=1), quantified by RT-qPCR targeting the HeV *N* gene (A) or TCID_50 a_ssay (B). HeV *N* gene expression was normalised to inoculum levels for each cell type. (C) Validation of FBL protein knockdown following transfection of 40 nM siRNA in TeloHAEC (120 h), HSAEC1-KT (120 h), NuLi-1 (72 h) and BE(2)-M17 cells (72 h). Results are representative of three separate biological replicates (n=3). *p<0.05 between inoculum and 24 hpi values according to two-tailed Student t test.

### FBL modulates HeV-g1 replication in a cell-dependent manner

The effect of FBL knockdown on HeV-g1 replication was assessed across the four cell lines. In HSAEC1-KT cells, a substantial reduction in HeV *N* mRNA was observed in siFBL cells compared to siNEG (Fig 2). At 24hpi, viral RNA synthesis was reduced 64% in siFBL cells, accompanied by a >1 log₁₀ decrease in infectious virus titres. The antiviral effect of siFBL was lost at 48 hpi, with viral RNA synthesis and virus titres comparable to siNEG. To determine whether alterations in viral replication were attributable to cytotoxic effects of siRNA treatment or virus infection, cell viability was assessed both in the absence and presence of viral infection (48 hpi). No significant reduction in cell viability was detected in siFBL cells, either in the absence or presence of infection. Cell viability was slightly elevated in siFBL cells infected with virus compared to siNEG. Use of siPLK1 as a positive control for apoptosis-induced cell death [20] confirmed assay sensitivity.

**Figure 2:**
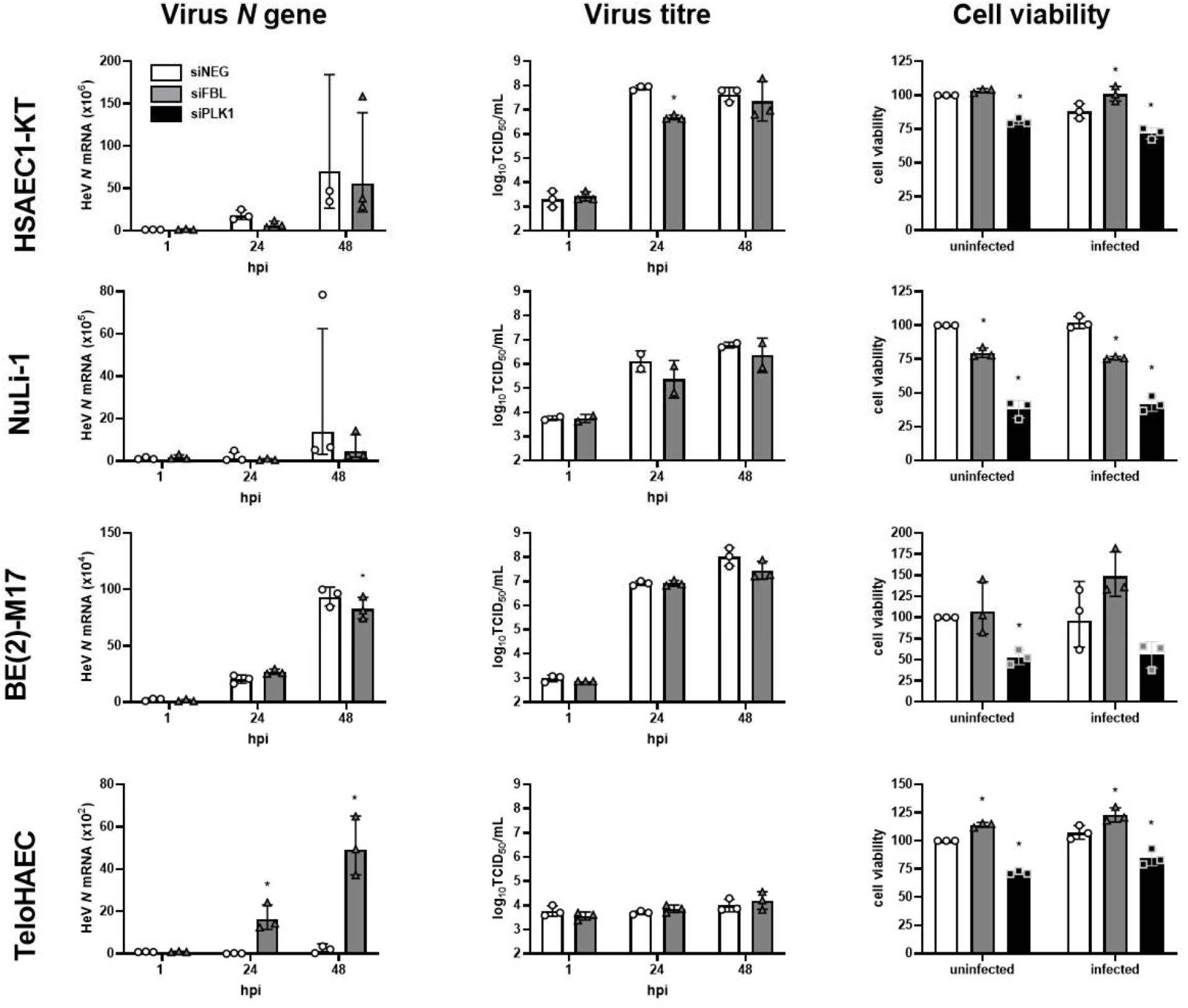
FBL modulates HeV-g1 replication in a cell-dependent manner. Cells were infected with HeV-g1 (TeloHAEC (MOI=1), HSAEC1-KT (MOI=0.2), Nuli-1 (MOI=1), BE(2)-M17 (MOI=0.05)). HeV *N* gene expression was normalised to inoculum levels for siNEG at 0 hpi. Cell viability was measured at 48 hpi. Results are representative of three separate biological replicates (n=3) *p<0.05 compared to siNEG according to two-tailed Student t test.

To investigate the impact of FBL knockdown on virus replication in other cell types, this experiment was extended to TeloHAEC, BE(2)-M17 and NuLi-1 cells. In NuLi-1 cells, FBL knockdown resulted in modest reductions in viral RNA synthesis that did not achieve statistical significance. A modest decrease in viral titre was also not significant. FBL knockdown caused a significant 20% reduction in cell viability in NuLi-1 cells, both in the absence and presence of infection. In BE(2)-M17 cells, FBL knockdown did not affect virus replication, virus titres or cell viability. Intriguingly, in TeloHAEC cells, HeV *N* gene synthesis was markedly increased in siFBL cells compared to siNEG, at both 24 and 48 hpi. Virus titres were unchanged and did not exceed baseline inoculum levels. Cell viability values were slightly increased in siFBL cells relative to siNEG, both in the absence and presence of infection.

### Differential modulation of henipavirus replication by FBL in HSAEC1-KT cells

We next investigated the effect of FBL knockdown on the replication of HeV-g2, NiV-B, NiV-M and CedV. HSAEC1-KT cells were used for these experiments given the robust HeV-g1 replication observed in these cells, as well as negligible impact of FBL knockdown on cell viability. Cells were infected with each henipavirus for 24 h, with virus replication measured by viral *N* gene quantification and TCID_50 a_nalysis (Supplementary Fig. 1). Robust replication of HeV-g2, NiV-B, NiV-M, and CedV were observed within 24 hpi, with *N* gene synthesis highest for HeV-g1 (>3,400,000-fold increase over inoculum), followed by NiV-B (>190,000-fold), NiV-M (>130,000-fold), CedV (>55,000-fold) and HeV-g2 (>200-fold). Increases in virus titres were highest for HeV-g1 (4.00 log_10T_CID_50 i_ncrease over inoculum), followed by NiV-M (3.42 log_10T_CID_50)_, NiV-B (3.07 log_10T_CID_50)_, HeV-g2 (2.74 log_10T_CID_50)_ and CedV (2.24 log_10T_CID_50)_. FBL knockdown resulted in significant reductions in *N* gene expression for HeV-g2 (58.2 ± 17.8% reduction standard deviation (SD)) and NiV-B (58.2 ± 29.4 (SD)%) at 24 hpi (Fig. 3), and modest, non-significant reductions for NiV-M (27.6 ± 49.8 (SD)%) and CedV (20.5 ± 27.2 (SD)%). No differences in virus titres were observed between siNEG and siFBL across the experiment.

**Figure 3:**
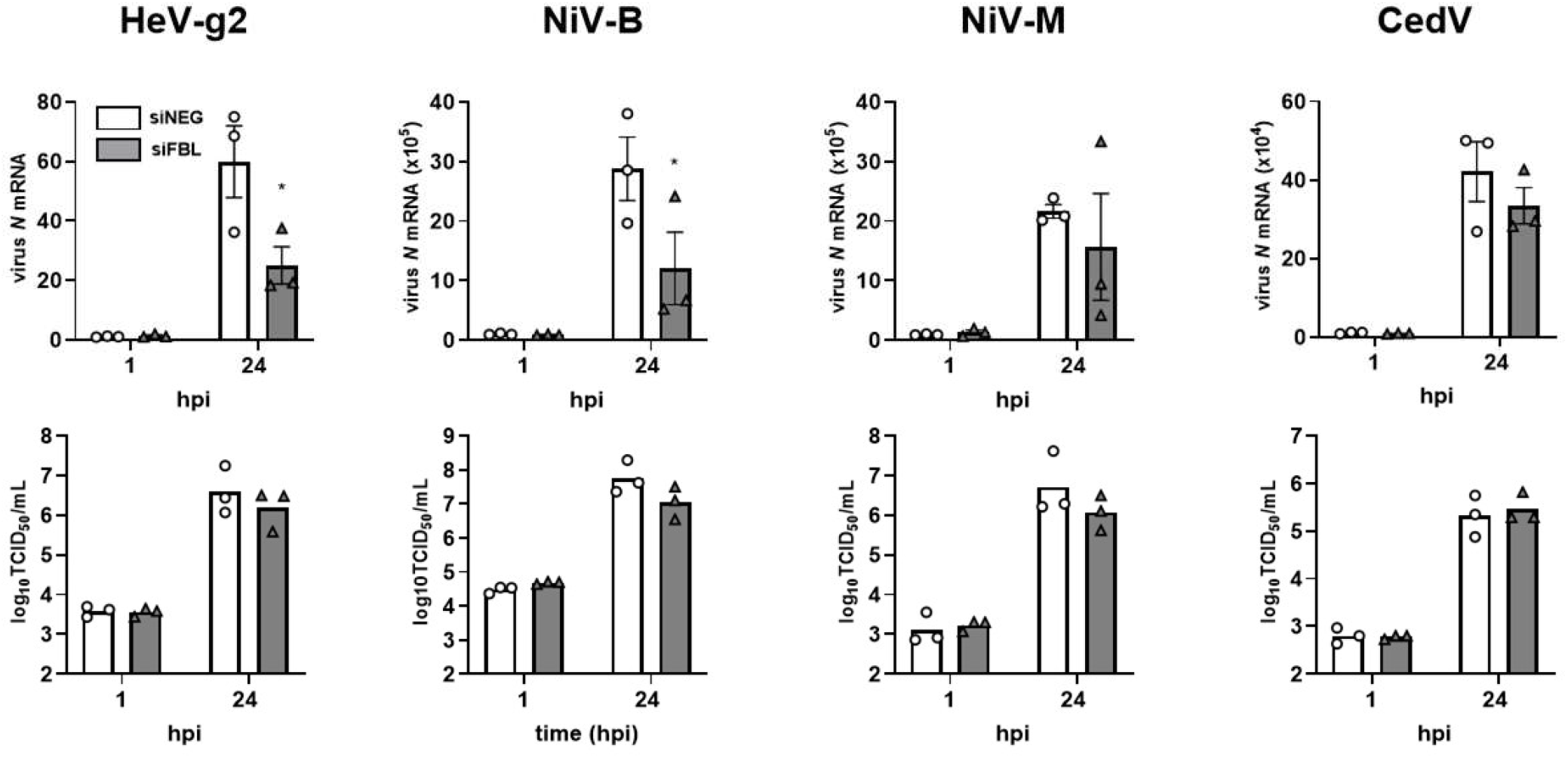
FBL differentially modulates henipavirus replication in HSAEC1-KT cells. HeV *N* gene expression was normalised to inoculum levels for siNEG at 0 hpi. Results are representative of three separate biological replicates. *\**p<0.05 compared to siNEG according to two-tailed Student t test.

### FBL depletion suppresses henipavirus transcription while amplifying innate immune and ISG responses

Transcriptomics was employed to investigate the impact of FBL knockdown on both viral and cellular gene expression during infection of HSAEC1-KT cells. Analysis of total virus transcript levels demonstrated a significant reduction between siNEG and siFBL at 24 hpi (Fig. 4A). A significant reduction in *N* and *M* gene reads in siFBL cells was observed (Fig. 4B, C), while *P*/*V*/*C*, and *G* transcripts were unaffected. *F* and *L* transcripts were significantly elevated in siFBL cells.

**Figure 4:**
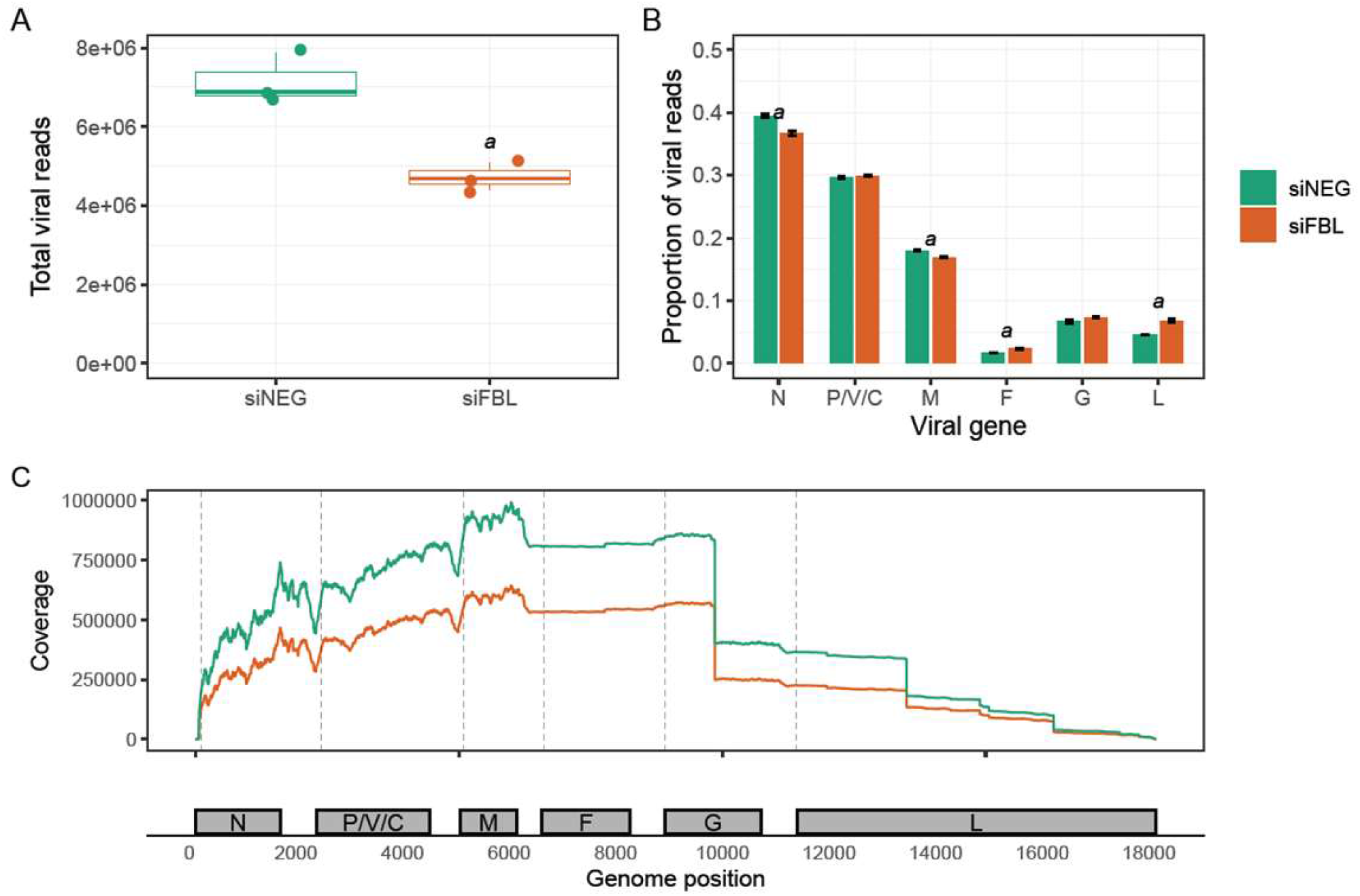
FBL depletion suppresses transcription of select HeV-g1 genes in HSAEC1-KT cells. (A) Total RNA-seq reads mapping to Hendra-g1 genome counted by the featureCounts programin cells treated with siNEG or siFBL, followed by HeV-g1 infection for 24 hpi (MOI 0.2). Boxplots show the distribution of viral reads across biological replicates, with individual replicate values overlaid as points. *a* < 0.05. (B) Relative distribution of viral RNA reads mapping to individual viral genes (*N*, *P/V/C*, *M*, *F*, *G*, and *L*) in siNEG- and siFBL-treated cells. Bars represent the mean proportion of total viral reads mapping to each gene, with error bars indicating standard error between three replicates. Genes marked with *a* differ significantly between treatments (a = p<0.05). (C) Genome-wide coverage profile of viral RNA sequencing reads across the viral genome in siNEG (green) and siFBL (orange) samples. Coverage is plotted as the number of reads mapped at each genomic position. Dashed vertical lines indicate gene boundaries, and the genomic organization is shown below the plot.

Differential expression (DE) analysis was performed to identify host transcriptional changes associated by FBL knockdown, both in the absence and presence of HeV-g1 infection. In uninfected cells, comparison of siNEG and siFBL treatments identified 4,330 DE host genes (absolute log_2 f_old change > 0.5 and FDR adjusted p-value < 0.05, Supplementary Table 1), indicating that FBL knockdown had a substantial impact on the host gene expression. Independently, HeV-g1 infection resulted in 2,038 DE genes in siNEG cells (Supplementary Table 2).

To specifically assess how FBL knockdown altered the host response to infection, we examined the interaction between FBL depletion and HeV-g1 infection. This analysis identified 1,343 GE genes whose expression was significantly influenced by the combined effects of both variables (Supplementary Table 3).

To facilitate biological interpretation of these transcriptional changes, Gene Set Enrichment Analysis (GSEA) was performed to identify significantly enriched pathways (FDR-adjusted p-value < 0.05). Leading-edge genes from each enriched pathway were then intersected with the corresponding DE gene sets. Pathways were considered to provide stronger evidence of biologically meaningful perturbation when 1) at least 50% of their leading-edge genes were differentially expressed, and 2) the median absolute log_2 f_old change of these DE genes exceeded 1 (Supplementary Figure 2). This approach prioritised pathways that were supported by both coordinated gene set enrichment and substantials changes in the expression of key contributing genes.

HeV-g1 infection in control cells induced innate antiviral and inflammatory pathways (Figure 5A), including interferon signalling, cytokine and chemokine signalling, and antigen processing and presentation. Additional enrichment of virus-host interaction pathways, such as RSV-host interactions, likely reflects shared host antiviral mechanisms captured by virus-specific Reactome annotations. In contrast, FBL knockdown (in the absence of infection) induced a broad cellular stress and homeostatic response (Figure 5B), with pathways involved in endoplasmic reticulum (ER) stress sensing (including activating transcription factor 6 (ATF6)-mediated ER chaperone response and inositol-requiring enzyme 1 / X-box binding protein 1 (IRE1α–XBP1) chaperone response), cellular stress responses (forkhead box O (FOXO)-mediated stress & metabolism, and response of eukaryotic translation initiation factor 2 alpha kinase 1 (EIF2AK1) (heme-regulated inhibitor, HRI) to heme deficiency), metabolic adaptation (sterol regulatory element-binding protein (SREBP)-regulated cholesterol & steroid biosynthesis), as well as altered cellular architecture (extracellular matrix organization, anchoring fibril formation, and epithelial differentiation & cornification) robustly enriched (Figure 5B).

**Figure 5.**
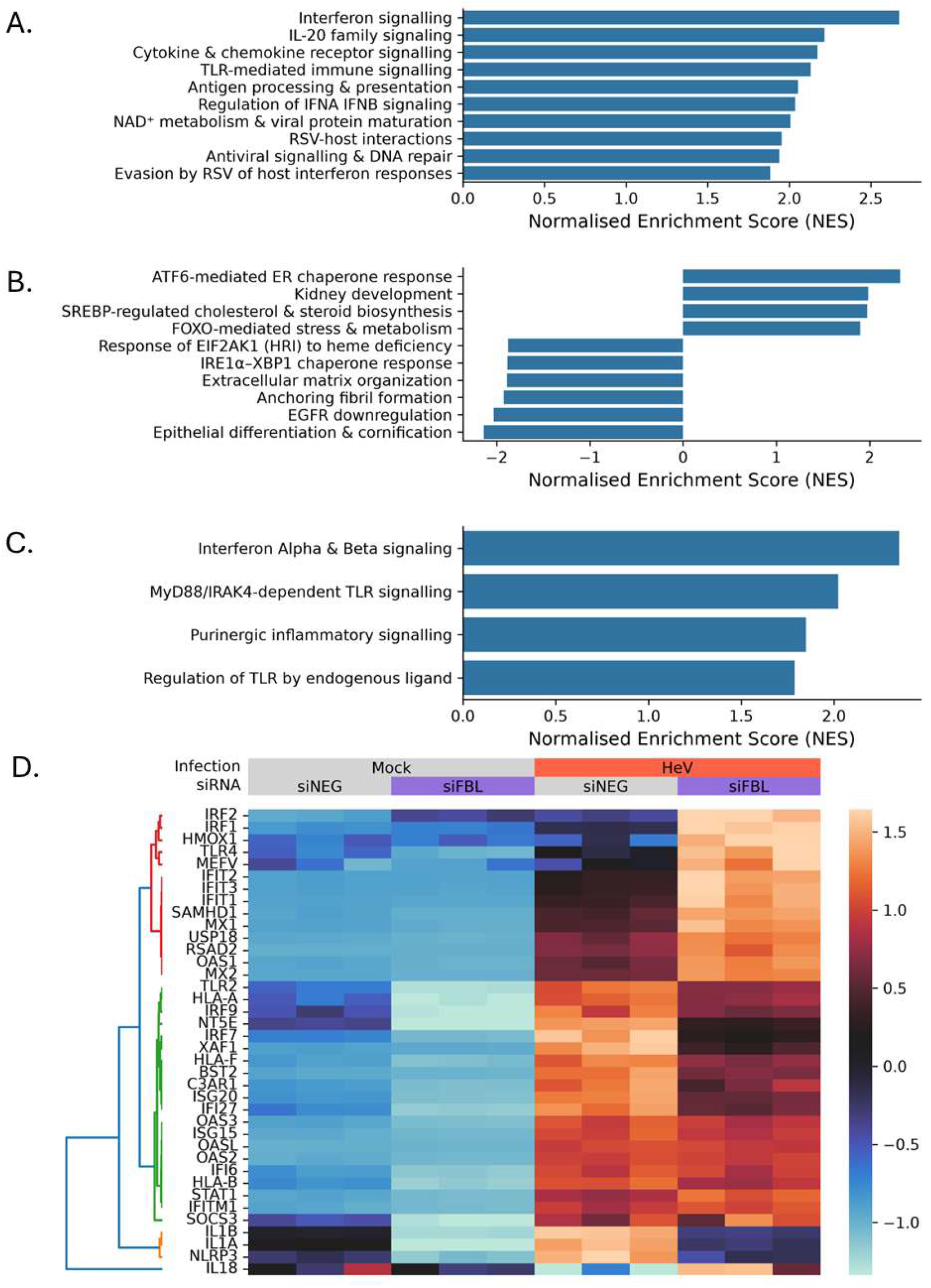
Innate immune and ISG responses to HeV-g1 are enhanced by FBL knockdown. (A-C) Barplots show the top 10 enriched pathways (sorted on FDR adjusted p-value) in (A) HeV-g1 infection, (B) siFBL and (C) siFBL and HeV-g1 infection interaction. Bars represent the normalized enrichment score (NES), with positive and negative values indicating enrichment towards the upregulated and downregulated ends of the DESeq2-ranked gene list, respectively. (D) Heatmap with hierarchical clustering dendrogram demonstrating the expression patterns of the DE leading edge genes from the pathways in (C). Gene expression data was transformed into z-scores. Samples (columns) are clustered based on experimental treatment groups, as indicated by the colored bars on top of the heatmap: infection (Mock – gray, HeV – red) and siRNA (siNEG – gray, siFBL – purple). Genes were clustered based on average correlation to identify expression patterns. Gene clusters (red/green/orange/blue) were defined with a correlation-distance threshold of 0.2.

**Figure 6:**
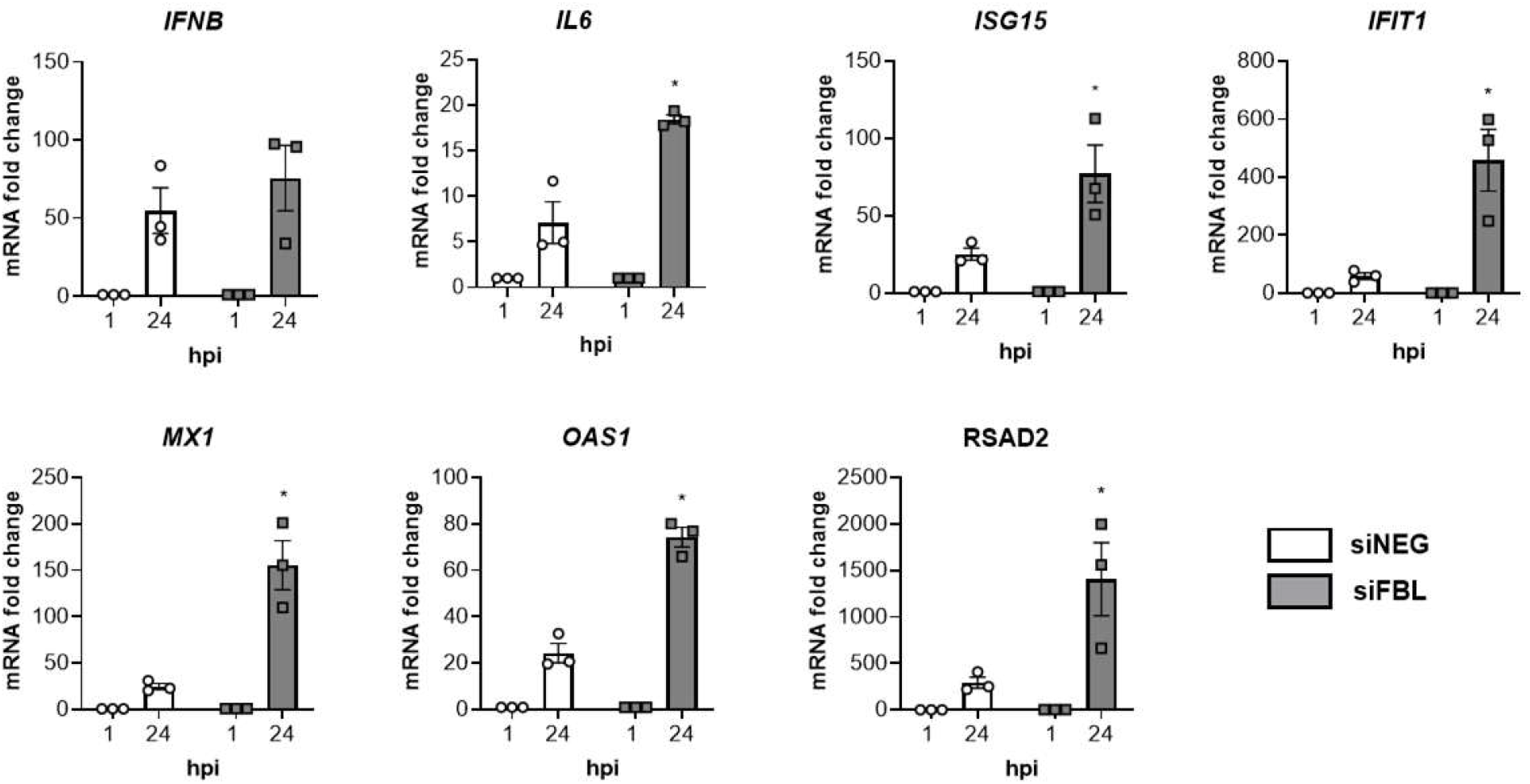
Validation of anti-viral immune signalling elevated by FBL knockdown in HSAEC1-KT cells. Host gene expression was measured by RT-qPCR in uninfected cells and those infected with HeV-g1 (MOI 0.2, 24 hpi). HeV *N* gene expression was normalised to inoculum levels for siNEG at 0 hpi. Results are reflective of three separate biological replicates (n=3) *p<0.05 in siFBL at 24 hpi compared to siNEG 24 hpi according to two-tailed Student t test.

Cellular pathways enriched in response to both FBL knockdown and HeV-g1 infection were associated with immune signalling, with interferon, toll-like receptor (TLR), and purinergic pathways highlighted (Figure 5C). Hierarchical clustering of differentially expressed leading-edge genes from these pathways identified distinct transcriptional response patterns, suggesting that the individual immune pathways exhibited divergent response to FBL depletion and HeV-g1 infection (Figure 5D). The expression of some genes, including *IRF1*, *IRF2*, *HMOX1*, *TLR4*, *MEFV*, *IFIT1*, *IFIT2*, *IFIT3*, *SAMHD1*, *MX1*, *MX2*, USP18, RSAD2, and OAS1 were enhanced during infection of siFBL cells relative to siNEG. Conversely, genes including *TLR2*, *HLA-A*, *IRF9*, *NT5E*, *IRF7*, *XAF1*, *HLA-F*, *BST2*, *C3AR1*, *ISG20*, and *IFI27* demonstrated a dampened response to HeV-g1 infection in siFBL cells (Figure 5D, green cluster). Furthermore, *IL1A*, *IL1B*, and *NLRP3* showed a response to HeV-g1 infection that was negated by FBL knockdown (Figure 5D, orange cluster). Interestingly, *IL18* clustered on its own, demonstrating an inverse expression pattern, with HeV-g1 infection reducing its expression in siNEG cells but increasing in siFBL cells (Figure 5D, blue cluster).

RT-qPCR analysis was performed in HSAEC1-KT cells to validate the host antiviral response factors identified by RNAseq. Consistent with the transcriptomic data, FBL knockdown resulted in a significant increase in the expression of *IL6*, *ISG15*, *IFIT1*, *MX1*, *OAS1* and *RSAD2* following HeV-g1 infection. This enhancement was observed in infected conditions, indicating that FBL knockdown amplifies virus-induced innate immune signalling rather than inducing basal immune activation.

## DISCUSSION

The highly pathogenic nature of HeV and NiV, coupled with evidence of human-to-human transmission and the current lack of medical countermeasures, highlights the need for antiviral strategies that can be rapidly deployed against multiple henipaviruses. HDA approaches seek to treat viral infections by targeting host factors that are essential for viral replication or contribute to disease severity, thereby providing a complementary alternative to conventional virus-targeted therapeutics. HDAs targeting cellular pathways required for virus replication may therefore retain activity against diverse viral strains while reducing the likelihood of viral escape. FBL emerged as a HDA candidate following a genome-wide siRNA screen in HeLa cells, where it was identified as one of the strongest host dependency factors for HeV-g1 and NiV-M replication [16].

A key objective of the present study was to determine whether the proviral role of FBL is conserved in cell types representative of tissues involved in henipavirus disease. Unlike HeLa cells, which harbour HPV-driven disruption of multiple cellular pathways [21], the HSAEC1-KT, NuLi-1 and TeloHAEC cell lines are immortalised derivatives of normal human airway epithelial and endothelial tissues. HSAEC1-KT cells were immortalised with hTERT and CDK4 [22], NuLi-1 cells with hTERT and HPV16 E6/E7 [23] and TeloHAEC cells with hTERT alone. Although BE(2)-M17 cells are a cancer-derived model, they provide a practical surrogate for neuronal cells, which are challenging to maintain in culture. Collectively, these models provide a physiologically relevant framework for examining host-virus interactions across the respiratory, endothelial and neurological tissues involved in henipavirus pathogenesis.

Our findings demonstrate that the contribution of FBL to henipavirus replication is influenced by cellular context. While FBL depletion reduced HeV-g1 replication in HSAEC1-KT cells and similarly impaired replication of HeV-g2 and NiV-B in this same cell type, equivalent effects were not observed in NuLi-1, TeloHAEC or BE(2)-M17 cells. Importantly, the antiviral phenotype identified in airway epithelial cells suggests that FBL may contribute to virus replication within selected cellular environments rather than across all susceptible tissues.

The differences observed between cell types may reflect variations in the biological functions of FBL within different tissues. FBL plays critical roles in ribosome biogenesis, translational regulation and cellular homeostasis [24]. The relative dependence of individual cell types on these processes is likely to differ, potentially influencing the extent to which viral replication relies upon FBL. The observation that only HSAEC1-KT cells displayed consistent antiviral responses following FBL knockdown suggests that either the requirement for FBL is greatest in these cells, or that FBL depletion more effectively engages antiviral restriction mechanisms within airway epithelial cells. Future studies comparing basal expression levels of FBL and associated ribosome biogenesis pathways across these cell types may help explain these differential phenotypes.

An alternative, but not mutually exclusive, explanation for the more modest antiviral effects observed here compared with the previous HeLa study is the degree of FBL knockdown achieved. Deffrasnes et al. reported near-complete depletion of FBL protein accompanied by profound inhibition of henipavirus replication, whereas knockdown efficiencies in the present study ranged between 60% and 80%. These findings may indicate a threshold requirement for FBL during henipavirus infection, whereby partial knockdown is sufficient to perturb replication in some cell types but not others. However, the differing outcomes between cell types despite comparable knockdown efficiencies suggests that biological differences between tissues may to contribute.

One of the most striking findings from this study was the association between FBL knockdown and enhanced innate immune signalling. Increased expression of multiple interferon-stimulated genes, including *IFIT1*, *IFIT2*, *IFIT3*, *MX1*, *MX2*, *OAS1*, *RSAD2* and *SAMHD1*, was observed upon infection of FBL knockdown cells. Notably, enhanced immune activation was primarily observed in infected cells rather than mock-infected controls, suggesting that FBL knockdown does not directly trigger antiviral signalling but instead amplifies virus-induced responses. Recent evidence provides a potential mechanistic explanation for this. FBL is responsible for generating 2’-O-methylated RNA species that act as endogenous antagonists of TLR7 and TLR8 signalling [25].

In addition, FBL-mediated RNA 2’-O-methylation has been shown to suppress innate immune sensing through MDA5-dependent type I interferon pathways, thereby facilitating viral infection in macrophages, while FBL depletion increases interferon-stimulated gene expression and restricts viral entry [26]. More recently, FBL knockdown was reported to inhibit respiratory syncytial virus (RSV) replication and enhance expression of multiple antiviral ISGs, further supporting a role for FBL as a negative regulator of antiviral immunity[27]. Together with our findings, these studies suggest that loss of FBL activity reduces the abundance of immunosuppressive RNA modifications, enhancing cellular responsiveness to RNA sensing pathways and amplifying virus-induced innate immune responses.

The viral transcriptomic data further suggest that FBL does not exert uniform effects across the henipavirus genome. While *N* and *M* transcript abundance was reduced following FBL depletion, transcripts corresponding to *F* and *L* were increased. This non-uniform response argues against a simple global suppression of viral RNA synthesis and instead suggests more complex effects on viral transcription, transcript stability, translation or host regulatory mechanisms. The biological significance of these gene-specific changes remains unclear but highlights that FBL likely influences multiple aspects of virus-host interactions beyond supporting replication alone.

Collectively, our findings refine the current understanding of FBL as a henipavirus host factor. Rather than functioning as a universally required proviral factor across all cell types, FBL appears to regulate infection in a tissue- and virus-dependent manner. The enhanced innate immune responses observed following FBL knockdown further support FBL as an underappreciated regulator of antiviral signalling. These observations support continued investigation of FBL biology during henipavirus infection while emphasising the importance of validating host-factor dependencies in biologically relevant cellular systems beyond the transformed cell lines commonly used for discovery screens.

## MATERIALS AND METHODS

### Cells and cell culture conditions

Telo-HAEC (ATCC CRL-4052), HSAEC1-KT (ATCC CRL-4050) and NuLi-1 (ATCC CRL-4011) cells were maintained in their respective Lonza bronchial epithelial growth media (CC-3202, CC-3118, and CC-3170). Telo-HAEC cells were cultured in CC-3202 supplemented with 2% of the supplied serum. BE(2)-M17 cells (ATCC CRL-2267) were maintained in MEM/F-12 (1:1; Gibco) supplemented with 2.5% fetal bovine serum. All cells were maintained at 37 °C with 5% CO₂ and 95% humidity and routinely confirmed to be mycoplasma-negative.

### Viruses

Infections involving HeV-g1 (Horse/1994/Hendra QLD/Australia), HeV-g2 (Horse/2015/Gympie QLD/Australia), NiV-M (Human/1999/Malaysia) and NiV-B (Human/2001/Bangladesh), and CedV (Flying Fox/2009/Cedar Grove QLD/Australia) were conducted at the CSIRO Australian Centre for Disease Preparedness at bio-safety level (BSL)-4. Cells were infected with virus at the MOI indicated for each cell line: MOI = 1 for Telo-HAEC and NuLi-1 cells, MOI = 0.2 for HSAEC1-KT cells, and MOI = 0.05 for BE(2)-M17 cells.

### siRNA transfection

Cells were reverse-transfected with 40 nM siRNA using DharmaFECT1 transfection reagent (Dharmacon), in accordance with the manufacturer’s guidelines. Non-targeting control siRNA (Dharmacon siGENOME Non-Targeting siRNA catalog number D-001210-02-05) or SMARTpool siRNAs targeting FBL (Dharmacon catalog number M-011269-00-0005), or PLK1 M-003290-01-0005 were used. Following 4 h incubation in transfection complexes, cells were replenished with fresh growth medium and incubated for a total of 72 h or 120 h as indicated, with media changes every 48 h for extended incubations.

### TCID₅₀ assay

Virus titres were determined by 50% tissue culture infectious dose (TCID₅₀) assay using Vero cells (ATCC CRL-81) maintained in MEM supplemented with 1% HEPES and 10% FBS. Samples were serially diluted and added in quadruplicate to Vero cells seeded in 96-well plates. Cytopathic effect was assessed five days post-infection, and TCID₅₀ values were calculated using the Reed and Muench method [28].

### Real-time quantitative PCR (RT-qPCR)

Total RNA was extracted using Tri Reagent (Sigma-Aldrich) according to the manufacturer’s instructions. Between 500 ng and 1 µg of total RNA was reverse-transcribed using the SuperScript III First-Strand Synthesis System (Thermo Fisher Scientific). Real-time quantitative PCR was performed using TaqMan Fast Universal PCR Master Mix (Thermo Fisher Scientific) and gene-specific TaqMan probes, including assays targeting viral nucleoprotein genes and host response genes (FBL (ThermoFisher Scientific TaqMan gene expression probe Hs01070449_m1, IL6 (Hs00174131_m1), ISG15 (Hs01921425_s1), IFNB1 (Hs01077958_s1), IFIT1 (Hs03027069_s1), MX1 (Hs00895608_m1), OAS1 (Hs00973635_m1), RSAD2 (Hs00369813_m1)) with GAPDH (Hs02786624_g1), HPRT1 (Hs02800695_m1), or 18S rRNA (Hs99999901_s1) used as reference genes. The N genes of HeV, NiV and CedV were detected using probes APU7CTE, APH6KCA and APKCEV7, respectively. Reactions were run on a QuantStudio real-time PCR system using the following cycling conditions: 95 °C for 20 s, followed by 40 cycles of 95 °C for 3 s and 60 °C for 30 s. Data were analysed using QuantStudio Design and Analysis software.

### Protein extraction and Western blotting

Cells were seeded into six-well plates and lysed post-transfection using ice-cold RIPA buffer (50 mM Tris pH 7.4, 150 mM NaCl, 1% IGEPAL, 0.5% sodium deoxycholate, 0.1% SDS) supplemented with protease inhibitor cocktail (Roche). Lysates were clarified by centrifugation at 16,000 × g for 10 min at 4 °C, and protein concentrations were determined using the Pierce BCA Protein Assay Kit (Thermo Fisher Scientific). Equal amounts of protein (16 µg) were resolved on 4–12% Bis-Tris NuPAGE gels using MES running buffer and transferred onto 0.2 µm nitrocellulose membranes using the Trans-Blot Turbo system (Bio-Rad). Membranes were blocked in 5% bovine serum albumin in PBS for 1 h and incubated overnight at 4 °C with primary antibodies against fibrillarin (Abcam ab4566; 1:500) or β-actin (Abcam ab8227; 1:10,000). After washing in PBST (PBS with 0.1% Tween-20), membranes were incubated with HRP-conjugated secondary antibodies for 1 h at room temperature. Signal detection was performed using Clarity Western ECL substrate (Bio-Rad) and imaged on a Bio-Rad ChemiDoc system. Band intensities were quantified using ImageJ software, normalised to β-actin, and expressed as fold-change relative to non-targeting siRNA-transfected controls.

### Cell viability assay

Cell viability was assessed using the CellTiter-Glo 2.0 luminescent assay (Promega). Cells were seeded into white-walled 96-well plates, transfected with siRNA, and infected as described above. At 24 h post-infection, assays were performed according to the manufacturer’s instructions, and luminescence was measured using a BioTek Synergy HT microplate reader.

### RNA sequencing

Total RNA was isolated from cells in TRIzol using the RNeasy kit (Qiagen) per the manufacturer’s instructions. The quality and quantity of RNA was assessed for all samples using an Agilent 2100 Bioanalyzer. RNA-seq was performed by the Australian Genome Research Facility (AGRF). Illumina TruSeq stranded mRNA libraries were prepared, followed by sequencing on an Illumina Novaseq-6000 for generation of 100-bp single-end reads. Between ∼15 and 20 million raw single-end reads were obtained per sample. Raw data were assessed for overall quality using fastqc v0.11.8 (http://www.bioinformatics.babraham.ac.uk/projects/fastqc/)

### Differential expression (DE) analysis

Genes with >=1 count per million in >=25% of samples were retained. Read normalization and differential expression was conducted using DESeq2 [29] using a factorial design with interaction term (∼ knockdown + infection + knockdown:infection) to investigate (1) the impact of siRNA knockdown or HeV infection alone, and (2) the effect on the infection response by siRNA knockdown. Genes with an FDR-adjusted p-value < 0.05 and an absolute log_2 f_old change > 0.5 were considered significant.

### Gene Set Enrichment Analysis (GSEA)

Genes were ranked using DESeq2’s Wald statistic and then pre-ranked GSEA was conducted using gseapy [30] with Reactome Pathways 2024 as the gene set, a minimum gene set threshold of 10, and 1000 permutations. The leading-edge genes of enriched pathways (FDR adjusted p-value < 0.05) were compared with identified DE genes; those pathways that had both >=50% of their leading-edge genes also DE and a median absolute log_2 f_old change of >=1 were considered to be high confidence and were retained. Jaccard similarity on leading edge overlap (threshold = 0.35) and Leiden clustering [31] on leading edge gene overlap were used to cluster together similar enriched pathways. Pathways were then collapsed and a representative pathway term chosen to reduce redundancy.

### Hierarchical clustering

Hierarchical clustering of genes was performed on z-score transformed reads using SciPy’s [32] cluster.hierarchy.linkage function, with average linkage and correlation distance. Gene clusters were defined by cutting the dendrogram at a correlation-distance threshold of 0.2 so the average correlation of the clustered genes is >=0.8.

### Statistical analysis

Statistical significance was determined using two-way analysis of variance (ANOVA) using data from three independent biological replicates. A *P* < 0.05 was considered statistically significant. Data were plotted as mean ± standard error of the mean (SEM) using GraphPad Prism software.

---

**Supplementary Figure 1:**
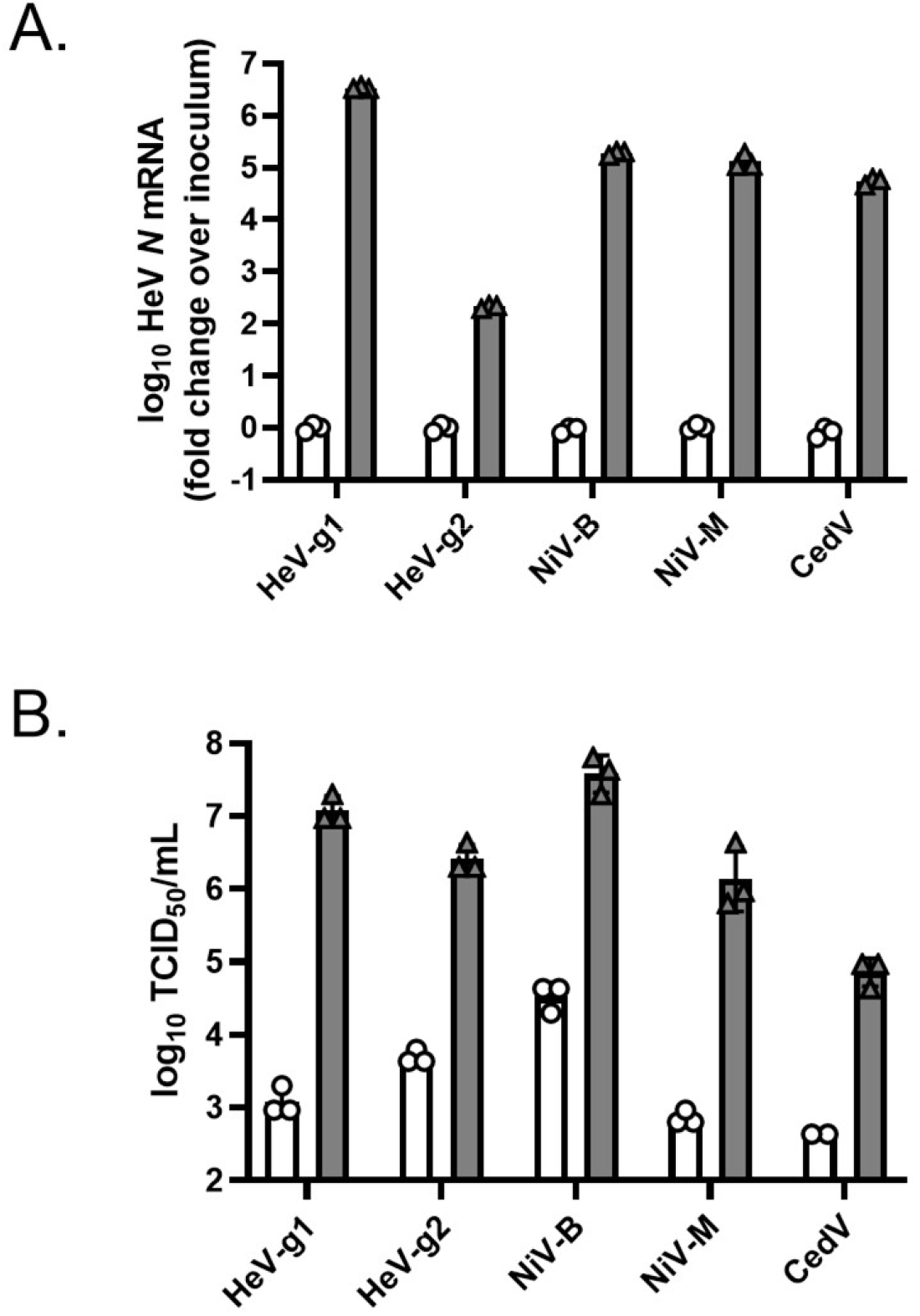
Comparative growth kinetics of henipaviruses in HSAEC1-KT cells. Henipavirus permissiveness was measured by RT-qPCR and infectious virus production was measured by TCID_50/_mL analyses. Inoculum samples were plotted as 1 hpi; n=3.

**Supplementary Figure 2.**
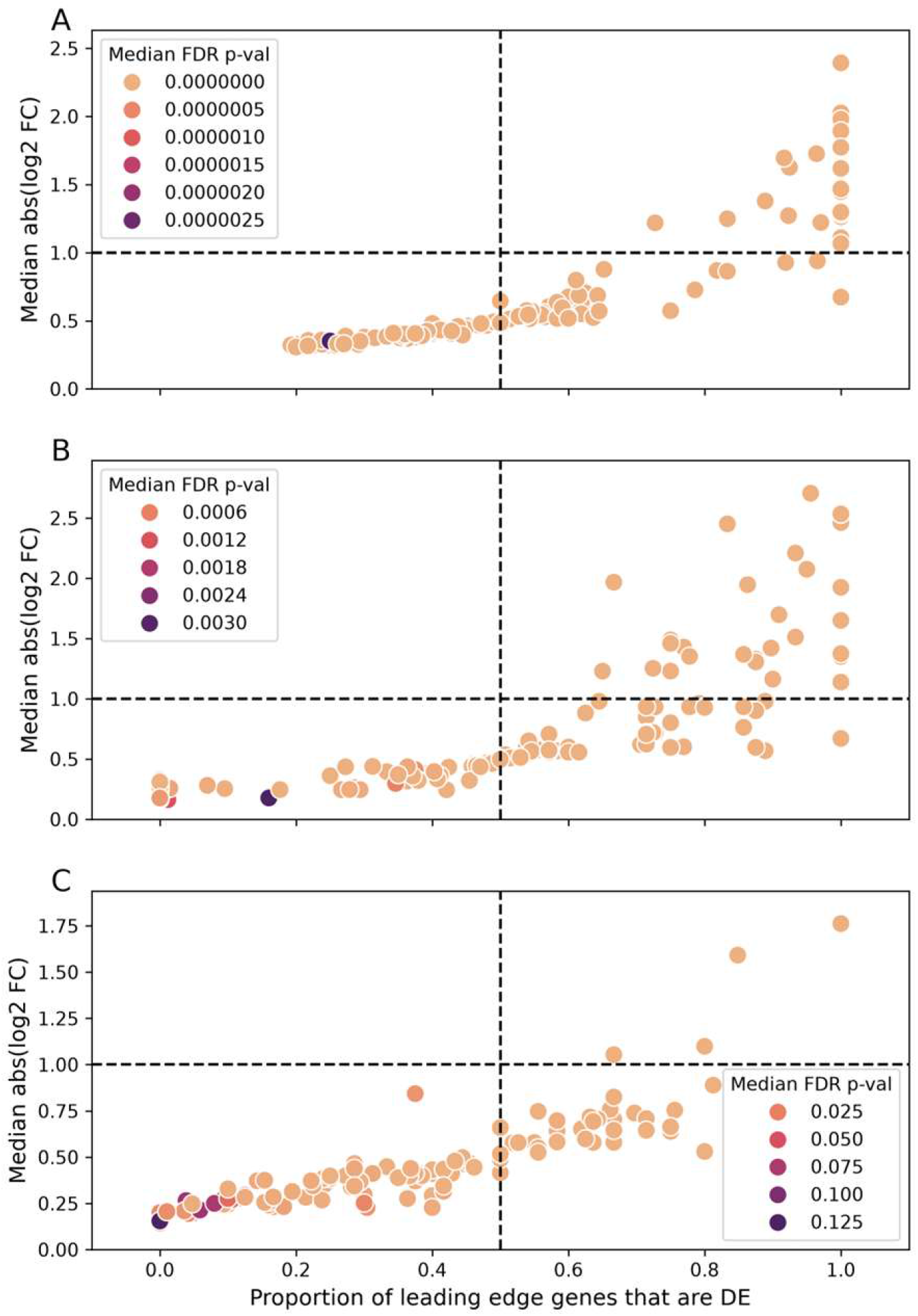
Enriched pathway leading edge genes – differentially expressed gene overlap. Scatterplots showing the proportion of leading-edge genes that were also DE (FDR adjusted-value < 0.05) for (A) HeV, (B) siFBL, and (C) HEV-siFBL enriched pathways. Each dot represents a single enriched pathway and the color represents the median FDR-adjusted p-value across all DE leading-edge genes. The dotted lines represent the threshold for both the proportion of DE leading-edge genes (0.5, vertical line) and median absolute log_2 f_old change (1, horizontal line). Those pathways in the top right-hand quadrant were considered high confidence.

## REFERENCES

1. Murray K, Selleck P, Hooper P, Hyatt A, Gould A, Gleeson L, et al. A morbillivirus that caused fatal disease in horses and humans. Science. 1995;268: 94–7. doi:10.1126/science.7701348

2. Chua KB, Bellini WJ, Rota PA, Harcourt BH, Tamin A, Lam SK, et al. Nipah virus: a recently emergent deadly paramyxovirus. Science. 2000;288: 1432–5. doi:10.1126/science.288.5470.1432

3. Khan S, Akbar SMF, Al Mahtab M, Uddin MN, Rashid MM, Yahiro T, et al. Twenty-five years of Nipah outbreaks in Southeast Asia: A persistent threat to global health. IJID regions. 2024;13: 100434. doi:10.1016/j.ijregi.2024.100434

4. Halpin K, Young PL, Field HE, Mackenzie JS. Isolation of Hendra virus from pteropid bats: a natural reservoir of Hendra virus. J Gen Virol. 2000;81: 1927– 1932. doi:10.1099/0022-1317-81-8-1927

5. Yob JM, Field H, Rashdi AM, Morrissy C, van der Heide B, Rota P, et al. Nipah virus infection in bats (order Chiroptera) in peninsular Malaysia. Emerg Infect Dis. 2001;7: 439–41. doi:10.3201/eid0703.010312

6. Chua KB, Goh KJ, Wong KT, Kamarulzaman A, Tan PS, Ksiazek TG, et al. Fatal encephalitis due to Nipah virus among pig-farmers in Malaysia. Lancet. 1999;354: 1257–9. doi:10.1016/S0140-6736(99)04299-3

7. O’Sullivan JD, Allworth AM, Paterson DL, Snow TM, Boots R, Gleeson LJ, et al. Fatal encephalitis due to novel paramyxovirus transmitted from horses. Lancet. 1997;349: 93–5. doi:10.1016/s0140-6736(96)06162-4

8. Gurley ES, Montgomery JM, Hossain MJ, Bell M, Azad AK, Islam MR, et al. Person-to-person transmission of Nipah virus in a Bangladeshi community. Emerg Infect Dis. 2007;13: 1031–7. doi:10.3201/eid1307.061128

9. Ukoaka BM, Okesanya OJ, Daniel FM, Ahmed MM, Udam NG, Wagwula PM, et al. Updated WHO list of emerging pathogens for a potential future pandemic: Implications for public health and global preparedness. Infez Med. 2024;32: 463–477. doi:10.53854/liim-3204-5

10. Middleton D, Pallister J, Klein R, Feng Y-R, Haining J, Arkinstall R, et al. Hendra virus vaccine, a one health approach to protecting horse, human, and environmental health. Emerg Infect Dis. 2014;20: 372–9. doi:10.3201/eid2003.131159

11. Pallister J, Middleton D, Wang L-F, Klein R, Haining J, Robinson R, et al. A recombinant Hendra virus G glycoprotein-based subunit vaccine protects ferrets from lethal Hendra virus challenge. Vaccine. 2011;29: 5623–30. doi:10.1016/j.vaccine.2011.06.015

12. Geisbert TW, Mire CE, Geisbert JB, Chan Y-P, Agans KN, Feldmann F, et al. Therapeutic treatment of Nipah virus infection in nonhuman primates with a neutralizing human monoclonal antibody. Sci Transl Med. 2014;6: 242ra82. doi:10.1126/scitranslmed.3008929

13. Bossart KN, Geisbert TW, Feldmann H, Zhu Z, Feldmann F, Geisbert JB, et al. A neutralizing human monoclonal antibody protects african green monkeys from hendra virus challenge. Sci Transl Med. 2011;3: 105ra103. doi:10.1126/scitranslmed.3002901

14. Bossart KN, Geisbert TW, Feldmann H, Zhu Z, Feldmann F, Geisbert JB, et al. A neutralizing human monoclonal antibody protects african green monkeys from hendra virus challenge. Sci Transl Med. 2011;3: 105ra103. doi:10.1126/scitranslmed.3002901

15. Playford EG, Munro T, Mahler SM, Elliott S, Gerometta M, Hoger KL, et al. Safety, tolerability, pharmacokinetics, and immunogenicity of a human monoclonal antibody targeting the G glycoprotein of henipaviruses in healthy adults: a first-in-human, randomised, controlled, phase 1 study. Lancet Infect Dis. 2020;20: 445–454. doi:10.1016/S1473-3099(19)30634-6

16. Deffrasnes C, Marsh GA, Foo CH, Rootes CL, Gould CM, Grusovin J, et al. Genome-wide siRNA Screening at Biosafety Level 4 Reveals a Crucial Role for Fibrillarin in Henipavirus Infection. PLoS Pathog. 2016;12: e1005478. doi:10.1371/journal.ppat.1005478

17. Escaffre O, Borisevich V, Rockx B. Pathogenesis of Hendra and Nipah virus infection in humans. J Infect Dev Ctries. 2013;7: 308–11. doi:10.3855/jidc.3648

18. Escaffre O, Borisevich V, Carmical JR, Prusak D, Prescott J, Feldmann H, et al. Henipavirus pathogenesis in human respiratory epithelial cells. J Virol. 2013;87: 3284–94. doi:10.1128/JVI.02576-12

19. Wong KT, Tan CT. Clinical and pathological manifestations of human henipavirus infection. Curr Top Microbiol Immunol. 2012;359: 95–104. doi:10.1007/82_2012_205

20. Liu X, Erikson RL. Polo-like kinase (Plk)1 depletion induces apoptosis in cancer cells. Proc Natl Acad Sci U S A. 2003;100: 5789–94. doi:10.1073/pnas.1031523100

21. Goodwin EC, DiMaio D. Repression of human papillomavirus oncogenes in HeLa cervical carcinoma cells causes the orderly reactivation of dormant tumor suppressor pathways. Proceedings of the National Academy of Sciences. 2000;97: 12513–12518. doi:10.1073/pnas.97.23.12513

22. Ramirez RD, Sheridan S, Girard L, Sato M, Kim Y, Pollack J, et al. Immortalization of Human Bronchial Epithelial Cells in the Absence of Viral Oncoproteins. Cancer Res. 2004;64: 9027–9034. doi:10.1158/0008-5472.CAN-04-3703

23. Zabner J, Karp P, Seiler M, Phillips SL, Mitchell CJ, Saavedra M, et al. Development of cystic fibrosis and noncystic fibrosis airway cell lines. American Journal of Physiology-Lung Cellular and Molecular Physiology. 2003;284: L844– L854. doi:10.1152/ajplung.00355.2002

24. Tollervey D, Lehtonen H, Jansen R, Kern H, Hurt EC. Temperature-sensitive mutations demonstrate roles for yeast fibrillarin in pre-rRNA processing, pre-rRNA methylation, and ribosome assembly. Cell. 1993;72: 443–457. doi:10.1016/0092-8674(93)90120-F

25. Alharbi AS, Sapkota S, Zhang Z, Jin R, Rupasinghe E, Jayasekara WSN, et al. 2′-O-Methyl-guanosine RNA fragments antagonize TLR7 and TLR8 to limit autoimmunity. Nat Immunol. 2026;27: 762–775. doi:10.1038/s41590-026-02429-2

26. Li P, Liu Y, Song R, Zhao L, Yang J, Lu F, et al. RNA 2’-O-Methyltransferase Fibrillarin Facilitates Virus Entry Into Macrophages Through Inhibiting Type I Interferon Response. Front Immunol. 2022;13. doi:10.3389/fimmu.2022.793582

27. Ulloa-Aguilar JM, Cruz-Holguin VJ, Benítez-Zeferino YR, Guzmán-Bautista ER, García-Cordero J, De Jesús-González LA, et al. Human respiratory syncytial virus regulates the expression of interferon-stimulated genes through modulation of fibrillarin. Front Cell Infect Microbiol. 2026;16. doi:10.3389/fcimb.2026.1706028

28. Reed LJ, Muench H. A SIMPLE METHOD OF ESTIMATING FIFTY PER CENT ENDPOINTS12. Am J Epidemiol. 1938;27: 493–497. doi:10.1093/oxfordjournals.aje.a118408

29. Love MI, Huber W, Anders S. Moderated estimation of fold change and dispersion for RNA-seq data with DESeq2. Genome Biol. 2014;15: 550. doi:10.1186/s13059-014-0550-8

30. Fang Z, Liu X, Peltz G. GSEApy: a comprehensive package for performing gene set enrichment analysis in Python. Bioinformatics. 2023;39. doi:10.1093/bioinformatics/btac757

31. Traag VA, Waltman L, van Eck NJ. From Louvain to Leiden: guaranteeing well-connected communities. Sci Rep. 2019;9: 5233. doi:10.1038/s41598-019-41695-z

32. Virtanen P, Gommers R, Oliphant TE, Haberland M, Reddy T, Cournapeau D, et al. SciPy 1.0: fundamental algorithms for scientific computing in Python. Nat Methods. 2020;17: 261–272. doi:10.1038/s41592-019-0686-2

